# Metabolically-versatile *Ca.* Thiodiazotropha symbionts of the deep-sea lucinid clam *Lucinoma kazani* have the genetic potential to fix nitrogen

**DOI:** 10.1101/2024.04.05.588213

**Authors:** Lina Ratinskaia, Stas Malavin, Tal Zvi-Kedem, Simina Vintila, Manuel Kleiner, Maxim Rubin-Blum

**Affiliations:** Biology Department, National Institute of Oceanography, Israel Oceanographic and Limnological Research (IOLR), Haifa, 3108000 Israel; Department of Marine Biology, Leon H. Charney School of Marine Sciences, University of Haifa, Haifa, 3498838 Israel; Zuckerberg Institute for Water Research, The Jacob Blaustein Institutes for Desert Research, Ben-Gurion University of the Negev, Sde Boker Campus 8499000, Israel; Department of Plant and Microbial Biology, North Carolina State University, Raleigh, NC, 27695, USA

## Abstract

Lucinid clams are one of the most diverse and widespread symbiont-bearing animal groups in both shallow and deep-sea chemosynthetic habitats. Lucicnids harbor *Ca*. Thiodiazotropha symbionts that can oxidize inorganic and organic substrates such as hydrogen sulfide and formate to gain energy. The interplay between these key metabolic functions, nutrient uptake and biotic interactions in *Ca*. Thiodiazotropha is not fully understood. We collected *Lucinoma kazani* individuals from next to a deep-sea brine pool in the eastern Mediterranean Sea, at a depth of 1150 m and used Oxford Nanopore and Illumina sequencing to obtain high-quality genomes of their *Ca.* Thiodiazotropha gloverae symbiont. The genomes served as the basis for transcriptomic and proteomic analyses to characterize the *in situ* gene expression, metabolism and physiology of the symbionts. We found genes needed for N_2_ fixation in the deep-sea symbiont’s genome, which, to date, were only found in shallow-water *Ca*. Thiodiazotropha. However, we did not detect the expression of these genes and thus the potential role of nitrogen fixation in this symbiosis remains to be determined. We also found the high expression of carbon fixation and sulfur oxidation genes, which indicates chemolithoautotrophy as the key physiology of *Ca*. Thiodiazotropha. However, we also detected the expression of pathways for using methanol and formate as energy sources. Our findings highlight the key traits these microbes maintain to support the nutrition of their hosts and interact with them.

## Introduction

Chemosynthesis, that is, the assimilation of inorganic carbon or methane into biomass using chemical energy rather than sunlight, is a key process driving productivity in some marine environments such as shallow water sediments and next to deep-sea vents and seeps [1]. Chemosynthetic bacteria and archaea established symbioses with an array of animal and protist hosts, allowing these organisms to thrive in marine habitats, such as hydrothermal vents, hydrocarbon seeps, deep organic falls and shallow-water seabed [2–4]. These associations differ in the taxonomic and functional diversity of the symbionts, the specificity of symbiont-host association, localization and transmission mechanisms [3, 5, 6].

Most symbionts are metabolic specialists, using a limited range of carbon and energy sources. Yet, some stand out exhibiting broader metabolic capacity. For example, the sulfur-oxidizing symbionts of bathymodioline mussels from hydrothermal vents can obtain energy from hydrogen [7] and can be mixotrophs, as they can use some organic carbon sources in addition to CO_2_ [8]. The bona fide chemosynthetic symbionts often co-occur with other microbes that can use small organic compounds as energy and carbon sources. For example, *Cycloclasticus* use short-chain alkanes in *Bathymodiolus heckerae* and *Methylophaga* methylotrophs sustain growth using methanol, a byproduct of methane oxidation [9, 10].

One remarkable example of chemosynthetic symbioses are lucinid (Lucinidae) clams and their sulfur-oxidizing symbionts. Lucinidae is a species-rich clade, usually colonizing shallow-water sediments in the vicinity of seagrass meadows and coral reefs, but some species are found in deep-sea chemosynthetic habitats, including in the vicinity of vents and seeps [11]. Lucinids are characterized by their obligate association with intracellular chemosynthetic bacteria, most often with *Ca.* Thiodiazotropha (Chromatiales, Sedimaenticolaceae), and rarely with other gammaproteobacteria, such as Thiohalomonadales species [12, 13]. Similar to other Chromatiales symbionts, such as those of tubeworms [14–17], *Ca.* Thiodiazotropha fuels carbon fixation via the Calvin-Benson-Bassham (CBB) cycle primarily using the oxidation of sulfide, as well as other inorganic sulfur compounds. Yet, their energy sources expand beyond sulfur and may include the oxidation of hydrogen, methanol and formate using electron acceptors such as oxygen and nitrate [12, 13, 18, 19]. Several *Ca.* Thiodiazotropha genotypes are diazotrophs that can fix dinitriogen [20, 21], however, such diazotrophic symbionts were only described for shallow water and warm habitats [13].

While shallow-water lucinids are widely studied, the research of deep-sea lucinids is limited, given the difficulty of deep-sea sample collection. Here we focus on the symbionts of the deep-sea lucinid species *Lucinoma kazani*, endemic to the Mediterranean Sea [22, 23]. *L. kazani* is a close relative of Codakiinae lucinid *Lucinoma borealis* [11]. The latter was described in the North Atlantic at depths down to 165 m and consistently hosts *Ca*. T. gloverae symbionts [13, 24]. While shallow-water *Ca*. T. gloverae can fix dinitrogen, those found in *L. borealis* from the cold deep-sea waters lacked the genetic potential for diazotrophy. In contrast, *L. kazani* is found in the deep eastern Mediterranean Sea, where the water is generally warm (∼14 °C) [25, 26]. Specifically, we found large populations of *L. kazani* near Palmahim Disturbance brine pools, which spew warm, ∼ 22 °C brines [27]. Asking whether the unique conditions in the deep eastern Mediterranean Sea can lead to unique adaptations of *L. kazani* symbionts, we assembled high-quality metagenome-assembled genomes (MAGs) using both long (Oxford Nanopore) and short (Illumina) reads, and assessed their functionality using genome-centered transcriptomics and proteomics.

## Materials and Methods

### Sample collection

In April 2021, we collected four *L. kazani* specimens that thrived on the sediment surface in a brine pool area in Palmahim Disturbance offshore Israel at a water depth of approximately 1150 m (32° 13’ 23.5" N 34° 10’ 42.19" E), using a SAAB Seaeye Leopard remotely operated vehicle (ROV) ’Yona’. The specimens were dissected onboard upon retrieval. Approximately 3 hours passed between scoop collection and processing (the scoop with animals was placed in ROV’s biobox), and gill tissue for consecutive DNA/RNA/protein extraction was preserved in RNAlater (4° for 24 hours, then RNAlater was decanted and specimens were kept at -80 °C). We recently showed that RNAlater is a good preservative for proteomic sample preservation in addition to the preservation of RNA/DNA [28].

### Stable isotope analyses

Tissue samples from five frozen clam specimens were lyophilized for 24 hours, homogenized and weighed before stable isotope analysis at Cornell University Stable Isotope Laboratory. The isotopic composition of organic carbon (n=5), nitrogen (n=5) and sulfur (n=4) was determined using a Thermo Delta V isotope ratio mass spectrometer (IRMS) interfaced to a NC2500 elemental analyzer (Sisma-Ventura et al., 2022). The isotopic composition of each sample was expressed as the relative difference between isotopic ratios in the sample and that in conventional standards (Vienna Pee Dee Belemnite, atmospheric N_2_ and Canyon Diablo Troilite for carbon, nitrogen and sulfur, respectively). Measured isotope ratios are reported in the δ per mille (‰) notation representing the deviation from the standards:

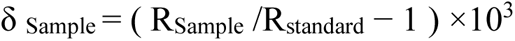

where, R = ^15^N/^14^N, ^13^C/^12^C or ^34^S/^32^S ratios.

### Preparation for omics

DNA, RNA and proteins were extracted from the four individuals using the AllPrep DNA/RNA/Protein Mini Kit (Cat. No. 80004, Qiagen). DNA libraries were constructed for these six individuals at HyLabs, Israel, and sequenced using 120 million 2x150 bp paired-end reads per sample with an Illumina NovaSeq at GENEWIZ, Germany. RNA libraries were constructed for these individuals at Novogene, Singapore, and sequenced using 100 million 2x150 bp paired-end reads per sample with an Illumina NovaSeq, following library construction with the NEBNext Ultra RNA Library Prep Kit for Illumina (Cat No. 7530) and ribosomal RNA depletion with the Ribo-Zero Plus rRNA Depletion Kit (Bacteria) (Cat No. 20037135) & Ribo-Zero Magnetic Kit (Plant Leaf). For long-read sequencing, DNA was extracted from an additional individual using DNeasy Blood & Tissue Kit (Qiagen). Long reads were produced by Minion (Oxford Nanopore), using R10.4.1 flow cells and SQK-NBD 114.24 Native Barcoding Kit 24 V14. Base calls from the generated raw Nanopore data were done using the super-accurate mode in the ONT Guppy base calling software version 6.4.2+97a7f06 with the dna_r10.4_e8.1_sup.cfg model. *Bioinformatics:* Host mitochondrial genomes were assembled and annotated with MitoZ [29] and the 16S and 18S rRNA gene sequences were extracted using phyloFlash v3.4.2 [30]. We carried out adapter trimming and error correction with tadpole.sh, using the BBtools suite following read preparation with the BBtools suite (Bushnell, B, sourceforge.net/projects/bbmap/). Metagenomes were assembled from short reads using SPAdes V3.15 with –meta k=21,33,66,99,127 parameters [31]. We performed downstream mapping and binning of metagenome-assembled genomes (MAGs) using DAStool, Vamb, Maxbin 2.0 and Metabat2 [32–35] within the Atlas V2.9.1 framework [36]. MAG quality was assessed using Checkm2 [37] and QUAST [38]. This analysis yielded a single high-quality, dereplicated symbiont MAG (99.98% completeness and 0.47% contamination, N50=57,715, 110 contigs). Long Naonopre reads (N50=1,614, longest read of 86,787 bases) were aligned to this genome using Minimap2 [39], and a hybrid genome was constructed from these aligned long reads and short reads, using the ‘isolate’ module in Nanophase 0.2.2 [40]. For functional annotation, we used the Rapid Annotations using the Subsystems Technology (RAST) server [41], and verified key annotations by BLAST search against the NCBI and UniProt databases. Prophage regions were identified with Phaster [42] and phage defense systems with DefenseFinder v1.2.0 [43].

RNA reads were quality-trimmed and mapped to the MAGs using BBmap with a 0.99 identity threshold (Bushnell, B, sourceforge.net/projects/bbmap/), and read counts were assigned to coding sequences using FeatureCounts [44]. The counts were normalized as transcripts per million (TPM) [45]. Phylogenomics was performed using the Codon Tree method in BV-BRC [46], which selects single-copy BV-BRC PGFams and analyzes single-copy genes using RAxML [47]. For selected features, genes/proteins were aligned with MAFFT [48], and treeing was performed with FastTree [49] and IQtree2 [50], using the best model.

### Protein extraction and peptide preparation for metaproteomics

We resuspended the protein precipitates from the four individuals extracted with the AllPrep DNA/RNA/Protein Mini Kit kit in 60 µl of SDT lysis buffer [4% (w/v) SDS, 100 mM Tris-HCl pH 7.6, 0.1 M DTT] and heated to 95°C for 10 min. The SDT protein mixture was cleaned up, reduced, alkylated and digested using the filter-aided sample preparation (FASP) protocol as described previously [51]. We performed all centrifugation steps mentioned below at 14,000 x g. We combined lysates (60 μl) with 400 μl of urea solution (8 M urea in 0.1 M Tris-HCl pH 8.5) and loaded it onto 10 kDa MWCO 500 μl centrifugal filters (VWR International) followed by centrifugation for 20 min. We washed filters once by applying 200 μl of urea solution followed by 20 min of centrifugation to remove any remaining SDS. We performed protein alkylation by adding 100 μl IAA solution (0.05 M iodoacetamide in urea solution) to each filter and incubating for 20 min at room temperature followed by centrifugation for 20 min. The filters were washed three times with 100 µL of urea solution with 15 min centrifugations, followed by a buffer exchange to ABC (50 mM ammonium bicarbonate). Buffer exchange was accomplished by three cycles of adding 100 μl of ABC buffer and centrifuging for 15 min. For tryptic digestion, we added 1 μg of MS grade trypsin (ThermoFisher Scientific) in 40 μl of ABC buffer to each filter and incubated for 16 hours in a wet chamber at 37°C. We eluted tryptic peptides by adding 50 μl 0.5 M NaCl and centrifuging for 20 min. Peptide concentrations were determined with the Pierce Micro BCA assay (ThermoFisher Scientific) following the manufacturer’s instructions.

*LC-MS/MS:* All proteomic samples were analyzed by 1D-LC-MS/MS as previously described [51] We loaded 1.2 μg of peptide from each sample onto a 5 mm, 300 µm ID C18 Acclaim® PepMap100 pre-column (Thermo Fisher Scientific) with an UltiMateTM 3000 RSLCnano Liquid Chromatograph (Thermo Fisher Scientific) in loading solvent A (2% acetonitrile, 0.05% trifluoroacetic acid). Elution and separation of peptides on the analytical column (75 cm x 75 µm EASY-Spray column packed with PepMap RSLC C18, 2 µm material, Thermo Fisher Scientific; heated to 60°C) was achieved at a flow rate of 300 nl min^-1^ using a 140 min gradient going from 95% buffer A (0.1% formic acid) and 5% buffer B (0.1% formic acid, 80% acetonitrile) to 31% buffer B in 102 min, then to 50% B in 18 min, and finally to 99% B in 1 min and ending with 99% . B. The analytical column was connected to a Q Exactive HF hybrid quadrupole-Orbitrap mass spectrometer (Thermo Fisher Scientific) via an Easy-Spray source. Eluting peptides were ionized via electrospray ionization (ESI). Carryover was reduced by a wash run (injection of 20 µl acetonitrile, 99% eluent B) between samples. MS1 spectra were acquired by performing a full MS scan at a resolution of 60,000 on a 380 to 1600 m/z window. MS2 spectra were acquired using data-dependent acquisition, selecting for fragmentation the 15 most abundant peptide ions (Top15) from the precursor MS1 spectra. A normalized collision energy of 25 was applied in the HCD cell to generate the peptide fragments for MS2 spectra. Other settings of the data-dependent acquisition included: a maximum injection time of 100 ms, a dynamic exclusion of 25 sec, and the exclusion of ions of +1 charge state from fragmentation. About 90,000 MS/MS spectra were acquired per sample.

### Protein identification and quantification

We constructed a protein sequence database for protein identification using the protein sequences predicted from the metagenome-assembled genomes obtained in this study. To identify peptides from the host, we used the annotated protein sequences of *Loripes orbiculatus*[52]. We added sequences of common laboratory contaminants by appending the cRAP protein sequence database (http://www.thegpm.org/crap/). The final database contained 101,483 protein sequences and is included in the PRIDE submission (see data access statement) in fasta format. Searches of the MS/MS spectra against this database were performed with the Sequest HT node in Proteome Discoverer 2.3.0.523 as previously described [53]. The peptide false discovery rate (FDR) was calculated using the Percolator node in Proteome Discoverer and only peptides identified at a 5% FDR were retained for protein identification. Proteins were inferred from peptide identifications using the Protein-FDR Validator node in Proteome Discoverer with a target FDR of 5%. To estimate species abundances based on proteinaceous biomass using the metaproteomic data we followed the previously described approach [54] with the added filter criterion of requiring two protein-unique peptides for a protein to be retained for biomass calculations.

## Results and Discussion

### Palmahim brine pool perimeter is inhabited by Lucinoma kazani

Clams formed a large, densely populated patch (several hundred square meters) near Palmahim brine pools (Figure 1). The clam shells were often covered with egg cases of *Galeus melastomus* sharks, which lay eggs in seeps likely due to the elevated temperatures that favor embryo development [55]. Thus, we assume that the temperature in shark/clam habitat is higher than the ambient 13.7 °C. In each scoop from this habitat, we found several live specimens among multiple empty shells.

**Figure 1:**
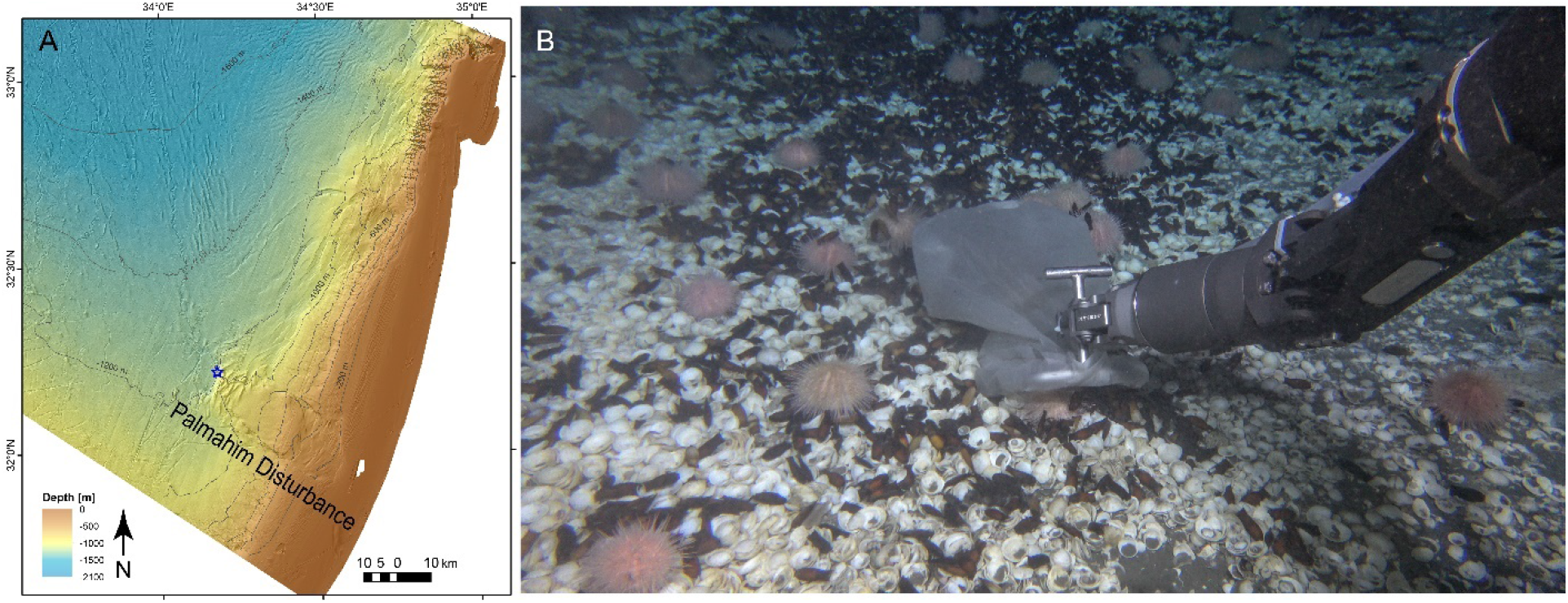
*Lucinoma kazani* inhabit the perimeter of deep-sea brine pools offshore Israel. A) The collection site location at the toe of Palmahim disturbance. B) Scoop collection of *L. kazani*.

Marker gene analysis indicated that the hosts were *Lucinoma kazani.* The mitochondrial *cytB* sequence was 99.15% similar to that of *L. kazani* (KF741674.1, the best hit in the NCBI). The 18S rRNA gene doesn’t distinguish between *L. kazani* and *L. borealis*, as the full-length nuclear 18S rRNA gene sequences in the four collected individuals were identical to that of *L. borealis* (AM774501.1, no *L. kazani* 18S rRNA gene sequences are available in the NCBI database).

We assembled a single symbiont 16S rRNA gene sequence from each metagenome using reads that mapped to the Silva138.1 16S rRNA gene sequence database. These sequences were identical among the four collected individuals, and 99.5% similar to those of the best hit in NCBI, *L. kazani* symbiont sequence AM236336.1. Phylogenomics showed that the *Ca.* Thiodiazotropha gloverae symbiont of *L. borealis* (GenBank: JAHXIE000000000.1, from Devon, United Kingdom) is the closest relative of *L. kazani* symbionts with 96.55% average nucleotide identity (**Supplementary Figure 1**). The symbionts were abundant in the gills of all four individuals (73-83% of peptide spectral matches).

Stable isotope analyses confirmed the nutritional dependence of *L. kazani* on the chemosynthetic symbionts. Bulk δ^13^C=-30.8±0.8‰ and δ^15^N=0.9±1.9‰ (n=5, **Supplementary Table ST1**) values were similar to δ^13^C=− 29.8‰ ± 0.9‰ and δ^15^N= − 2.5‰ ± 0.6‰ (n = 3) identified in the symbiont-bearing gills of *L. capensis* from the Namibian Shelf [56]. The negative δ^34^S values between − 18.9‰ and -1.6‰ indicated that chemosynthesis is driven by sulfur oxidation [57].

### Hybrid assembly improved the contiguity of the symbiont genome and identified large adhesin-like proteins

The hybrid assembly resulted in a more contiguous ∼4.9 Mb-long MAG consisting of 44 contigs with an N50=286,621 bp and the largest contig of 532,579 bp (**Table 1**). CheckM2 estimated completeness and contamination at 98.9% and 0.4%, respectively. Most importantly, the long reads were able to resolve long sequences (13,449 and 10,077 bp in length ) encoding several FhaB (filamentous hemagglutinin, COG3210) domains of adhesins (**Supplementary Figure S2**). These proteins are usually secreted to outer bacterial membranes and serve as attachment factors to host cells[58], playing a role in pathogenicity [59] and symbioses [60]. *L. kazani* symbionts indeed may secrete these adhesin-like proteins, as we identified several genes encoding secretion-related features in the neighborhood of the two adhesin-like protein-encoding sequences (e.g, those encoding proteins with HylD/AcrA-like domains, two-component system response regulator and sensor histidine kinase, type I secretion system permease/ATPase, **Supplementary Figure S2**). Apart from FhaB, the putative adhesins comprised additional eukaryote-like domains, often resembling RTX/MARTX toxins [61, 62], which may be involved in interactions of symbionts and their hosts [63, 64], facilitating not only adhesion but also cell surface cleavage [65]. These domains comprised the toxin-related Ca_2_^+^-binding protein (COG2931) and choice-of-anchor K domains, cadherin-like and bacterial Ig domains, as well as M36-family metallopeptidases, among others (**Supplementary Figure S2**). BLASTing against the NCBI database identified proteins with highly similar domains among *Ca.* Thiodiazotropha symbionts of various lucinids. Examples include the 5907 aa protein in *Ca.* Thiodiazotropha taylori (MBT3016762.1) and a 5144 aa protein in *Ca*. Thiodiazotropha ex. *Troendleina suluensis* (MCU7840510.1). As full-length sequences are often missing in MAGs produced by the short-read assemblies, the overview of adhesin diversity in *Ca*. Thiodiazotropha is incomplete. Similar adhesin-like proteins were highly expressed by symbionts of *Phacoides pectinatus* lucinids [66]. We, however, observed a limited expression of these sequences at the RNA level (no proteins found), hinting that the respective genes may be downregulated in established symbioses, or expressed by a small fraction of the symbiont population. Given the key role of adhesins in interactions between metazoans and bacteria, as well as their reoccurrence and expression in *Ca*. Thiodiazotropha, we hypothesize that these large proteins play a role in host colonization.

**Table 1:** Assembly statistics for metagenome-assembled genomes of *Lucinoma kazani* symbionts.

|  | Short-read assembly | Hybrid assembly |
| --- | --- | --- |
| Length (bp) | 4,346,866 | 4,873,710 |
| Contigs | 110 | 44 |
| N50 (bp) | 57,715 | 286,621 |
| Longest contig (bp) | 202,093 | 532,579 |
| Completeness (%) | 99.98 | 98.81 |
| Contamination (%) | 0.47 | 0.44 |

### Ca. T. gloverae ex L. kazani are metabolically versatile

The core, most well-expressed metabolic traits in *Ca.* T. gloveare ex. *L. kazani* included those involved in carbon fixation via the Calvin-Benson-Bassham (CBB) cycle and sulfur oxidation via the reverse dissimilatory sulfite reduction (DSR) pathway (**Figure 2, Supplementary Table ST2**). Similar to other Chromatiales symbionts [67, 68], *Ca.* T. gloverae ex. *L. kazani* uses the energy-efficient CBB pathway, driven by an H^+^-PPase and a closely coupled PPi-dependent 6-phosphofructokinase (PPi-PFK). The *pfP* and *hppA* genes that encode these features are neighbors that appear to be co-expressed and have high expression values (**Figure 2, Supplementary Table ST2**). The symbiont encoded two forms of ribulose bisphosphate carboxylase/oxygenase (RubisCO), which differ in their affinity to oxygen and CO_2_: form I has have high specificity factor, that is, functions better under low CO_2_ and high O_2_ conditions and form II enzymes, with low specificity factor [69–72]. This may reflect an adaptation to fluctuations in CO_2_ and O_2_ availability within the host’s bacteriocyte. Form I RubisCO was highly expressed compared to form II (11-79 more RNA reads mapped, and 8-14 times more protein spectra), suggesting that either oxygen was not limiting, or CO_2_ starvation. Other parts of the CBB pathway were also highly expressed. The CBB pathway is linked to glycogen storage, based on the substantial expression of the *glgABC* gene cluster. The symbiont is capable of gaining energy from the breakdown of these stored compounds, encoding and expressing a complete tricarboxylic acid cycle. We also found that several TRAP transporters were expressed, hinting at uptake of some organics, and thus mixotrophy.

**Figure 2:**
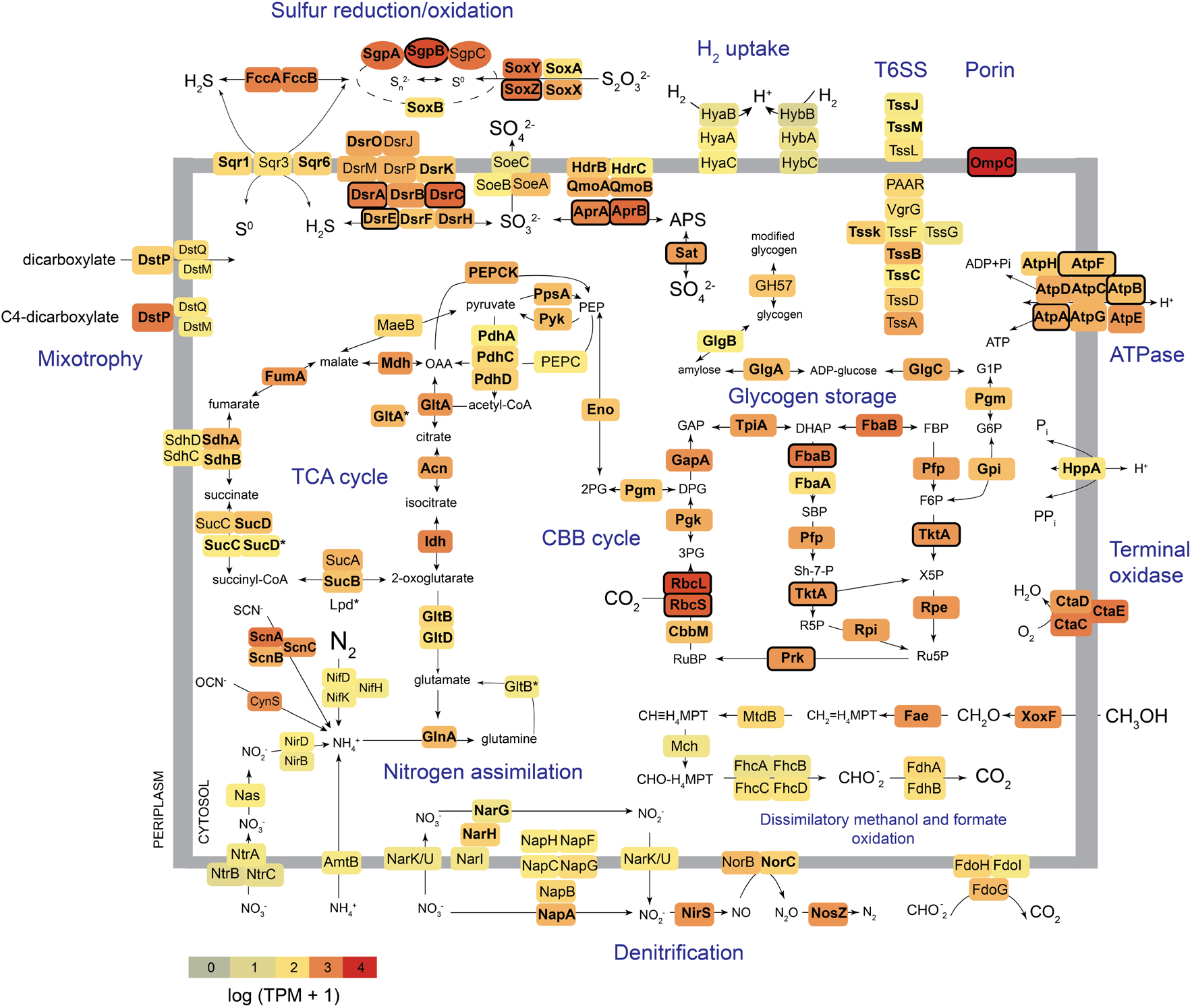
The key metabolic features in *Ca.* Thiodizaotropha gloverae ex. *Lucinoma kazani*. Color code is according to transcript abundance. Features detected at the protein level are marked in bold font. Those that constitute the top 50 most abundant proteins are framed. See Supplementary Table ST3 for protein annotations. TCA (tricarboxylic acid, Krebs) and CBB (Calvin-Benson-Bassham) cycles are abbreviated. * more than one gene copy present. ** The bona fide Lpd subunit of the 2-oxoglutarate dehydrogenase was not found, yet several variants of pyruvate dehydrogenase-associated lipoamide dehydrogenase were present, and may potentially complement the missing subunit.

The key enzymes involved in dissimilatory sulfide oxidation, including the adenylyl-sulfate (APS) reductase (AprAB), the sirohaem dissimilatory sulfite reductase (DSR) complex and sulfate adenylyltransferase (Sat), were detected as abundantly expressed at both the RNA and protein levels. Apart from Sat, the symbiont encoded the membrane-bound sulfite dehydrogenase (SoeABC). These two enzymes appear to synergistically catalyze the oxidation of sulfite in the purple sulfur bacterium *Allochromatium vinosum* [73]. Our expression analyses highlighted the important role of sulfur globule storage [74], as three types of sulfur globule proteins were highly expressed. In particular, SgpB was among the most well-expressed genes and most abundant proteins. The expression of the Sox system (SoxBXA and SoxYZ), needed to convert thiosulfate to sulfur globules, as well as the flavocytochrome c sulfide dehydrogenase FccAB, oxidizing sulfide to elemental sulfur [75], was substantial. We found three forms of sulfide: quinone reductases (Sqr1, 3 and 6), which differ in their affinity to sulfide [76], indicating the adaptability of the symbiont to unstable substrate concentrations. In summary, *Ca.* T. gloveare ex. *L. kazani* has a flexible rDSR pathway, typical of Chromatiales symbionts, such as those of siboglinid tubeworms [3].

*Lucinoma* symbionts use oxygen as a terminal electron acceptor but can reduce nitrate in the presence of oxygen [77]. In line with this observation, the expression of the CtaCDE terminal oxidase was high, yet the complete denitrification pathway was encoded and expressed. The symbionts may catalyze nitrate reduction using both low-affinity dissimilatory Nar and high-affinity periplasmic Nap nitrate reductases. Nap was expressed higher than Nar (also markedly higher at the protein level), potentially indicating nitrate limitation of denitrification [78, 79].

Unlike other *Ca.* T. gloverae [13], *L. kazani* symbiont encoded the canonical pathway of nitrate assimilation via Nas and NirBD. Yet, we found that the expression of assimilatory nitrate and nitrite reduction, as well as that of ammonium uptake, was generally low. In turn, both glutamine synthetase (GlnA) and glutamate synthase (GltBD) were moderately expressed. The very low expression of P-II nitrogen regulatory protein (GlnK) may indicate that ammonium is not limiting *Ca.* T. gloverae ex. *L. kazani* growth [80], which is likely given the millimolar levels of ammonium in Palmahim brines [27]. Additional ammonium can be derived from cyanate and thiocyanate, given that both cyanate hydratase and thiocyanate hydrolase were encoded and moderately transcribed by the *L. kazani* symbiont (**Figure 2, Supplementary Table ST2**).

While N2 fixation was attributed to date only to the symbionts of lucinids from shallow habitats [13], we found a complete cluster of genes involved in nitrogen fixation, in particular, those that encode the core subunits of molybdenum-iron nitrogenase (NifDK) and nitrogenase reductase and maturation protein NifH. The phylogeny of these genes is not congruent with *Ca.* Thiodiazotropha phylogenomics (**Figure 3, Supplementary Figures 1** and **3**). Most strikingly, the best BLAST hits of the key *nifD* (98.4% identity) and *nifK* (96.4% identity) were most similar to the genes from the distant clade hosted by *Notomyrtea botanica* (Notomyrt1 species). Hits to *nifD* genes from other *Ca*. Thiodiazotropha were below 95 and 94%, respectively (**Figure 3**). This hints at an evolutionary history of the nitrogenase cluster in this clade, with widespread gene losses, and potential secondary gains, which, however, are constrained within *Sedimenticolaceae*. Given that nitrogen fixation in lucinid symbionts is linked to shallow and warm habitats[13], temperature may play a role in the selection of this trait. Lucinid habitat in the deep easter Mediterranean Sea is characterized by aberrantly warm temperatures of ∼14 °C of the deep water column, however, the actual temperature may be even higher, given the inflow of ∼22 °C-warm brines. This may have played a role in retaining diazotrophy potential. Yet, we detected minimal expression values for this cluster at the RNA level, and no protein-level expression was found. Thus, nitrogen fixation may be limited in the host-associated *Thiodiazotropha*, for which other not as energetically-costly nitrogen sources such as ammonium and nitrate are available.

**Figure 3:**
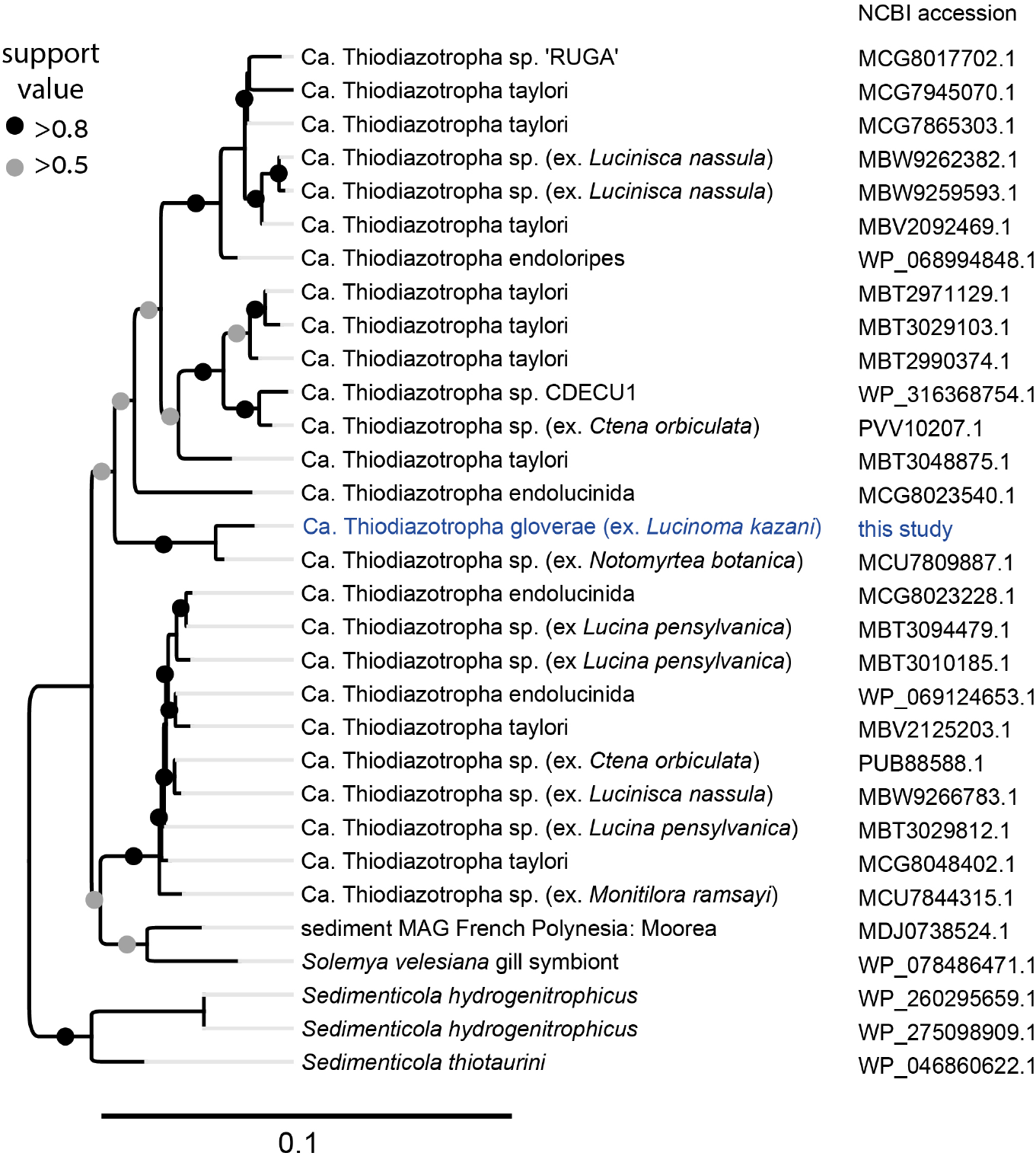
Phylogeny of 31 NifD amino acid sequences in the Thiodiazotropha clade. The maximum likelihood tree was constructed in FastTree, using the LG model. The tree was force-rooted at the *Sedimenticola* genus branch. The tree scale represents the number of substitutions per site. See Supplementary Figure S3 for an extended nucleotide tree.

Following previous studies [12, 13, 18, 19], our data suggests that the use of methanol, formate and hydrogen may support the energy metabolism in *Ca.* T. gloverae ex. *L. kazani*. The symbionts can conserve energy by oxidizing methanol to formaldehyde, formate and CO_2_, based on a gene cluster that encodes the lanthanide-dependent methanol dehydrogenase (XoxF), formaldehyde detoxification via the H4MPT pathway and the tungsten-containing formate dehydrogenase. Methanol appears to be a source of energy, but not carbon, as the key genes in formaldehyde assimilation pathways were not found. These include genes encoding the key enzymes of ribulose monophosphate pathway 3-hexulose-6-phosphate synthase (*rmpA*) and 3-hexulose-6-phosphate isomerase (*rmpB*), as well as in serine pathway, such *sgaA*, encoding serine--glyoxylate aminotransferase. The symbiont also encoded and expressed the respiratory membrane-bound formate dehydrogenase FdoGHI, which may allow the conservation of energy under anaerobic conditions using nitrate as an electron acceptor [81].

We identified genomic clues for the use of dihydrogen by *L. kazani* symbionts. Similar to their relative *Sedimenticola hydrogenitrophicus* that can grow hydrogenotrophically [82], the symbiont genome contained a cluster of genes encoding the HyaABC subunits of the H2-uptake [NiFe] hydrogenase, the accessory proteins, and hydrogen-sensing hydrogenase HoxCB. The *hyaA* and *hyaB* genes were present in two copies. The expression of these genes was low, indicating that similar to diazotrophy, host-associated symbionts were likely not using hydrogen under the *in situ* conditions at the time of sampling. Moreover, these hydrogenases can be directly linked to dinitrogen fixation, supplying electrons to sustain this process, as described in the purple bacterium *Rhodopseudomonas palustris* [83].

### Potential features responsible for biotic interactions of Ca. T. gloverae ex. L. kazani

We observed traits that can mediate the interaction between *Ca.* T. gloverae ex*. L. kazani* with the host, or other bacteria/competing strains. Similar to the symbionts of tubeworms, the most well-expressed gene/protein was an OmpC superfamily porin [16, 84]. These prior studies suggested that such porins may be crucial for maintaining stable symbiotic associations. Together with adhesin-like proteins, these porins may play a role in maintaining the symbioses, yet their role has not been experimentally tested so far.

The symbionts encoded and expressed components of the type VI secretion system (T6SS), which facilitates virulence, host immunomodulation and bactericidal activity in many bacterial lineages [85–87], and were previously found to be expressed by lucinid symbionts [18, 19]. These contact-dependent interbacterial “weapons” [88] are most likely to be involved in interactions with competing symbiont genotypes or cheaters during colonization of the host [18, 84, 89]. For example, in bobtail squid symbionts *Vibrio fischeri*, T6SS was shown to facilitate strain incompatibility, eliminating closely related competitor strains [89, 90].

We identified only a few phage-defense systems in the genome, while no CRISPR-Cas systems were found. These defense systems included LmuB SMC Cap4 nuclease II and LmuA effector hydrolase from a Lamassu family [91], and very few restriction-modification and toxin-antitoxin systems (e.g., sanaTA, [92]), often integrated into poorly-expressed regions identified as partial inactive prophages (**Supplementary Table ST2**). This suggests a limited phage predation pressure on the *L. kazani* symbionts.

## Conclusions

*Lucinoma kazani*, and their *Ca.* Thiodizotropha gloverae symbionts thrive in the warm deep-sea chemosynthetic habitat near the Mediterranean Sea brine pools. These conditions may determine their metabolic adaptations, for example, retaining the ability to fix dinitrogen. These deep-sea clam symbionts appear to maintain a versatile metabolism, modulating it via gene expression. For example, the preferential expression of cyanate/thiocyanate module over ammonium uptake, or changes in the expression of Sqr proteins with different affinities to sulfide. Further exploration of expression patterns under different environmental scenarios is needed to shed light on the interplay between different physiological strategies. Moreover, the heterogeneity of expression at a single symbiont level remains to be explored.

These genomes encode and express several features that can be involved in biotic interactions, including porins, secretions systems and adhesin-like proteins. We speculate that this versatility requires the maintenance of large genomes of ∼5 Mb, which is larger than those previously suggested for lucinid symbionts from extreme environments, such as ∼3 Mb genomes of *Ca*. T. “Aeq1” clade [13]. The discovery of key features such as toxin-like proteins in these large genomes was possible due to hybrid assemblies based on long-read sequencing.

## Data availability

The raw metagenomic reads, as well as the final MAGs, were submitted to NCBI in PRJNA1078112. The mass spectrometry proteomics data have been deposited to the ProteomeXchange Consortium via the PRIDE [93] partner repository with the dataset identifier PXD051186.

## Author contributions

MR-B, LR and MK conceived this study and acquired funding. LR, TZ-K and SM performed DNA/RNA/protein extraction and data analyses. SV and MK conducted proteomics. LR, MR-B, and MK wrote the paper with the contributions of all co-authors.

## Acknowledgments

The authors thank all individuals who helped during the expeditions, including onboard technical and scientific personnel, and the captains and crew of the E/Vs Bat Galim. We thank Ben Herzberg, Samuel Cohen, who operated Yona ROV. All LC-MS/MS measurements were made in the Molecular Education, Technology, and Research Innovation Center (METRIC) at North Carolina State University.

## Supplementary Material

**Supplementary Figure S1:**
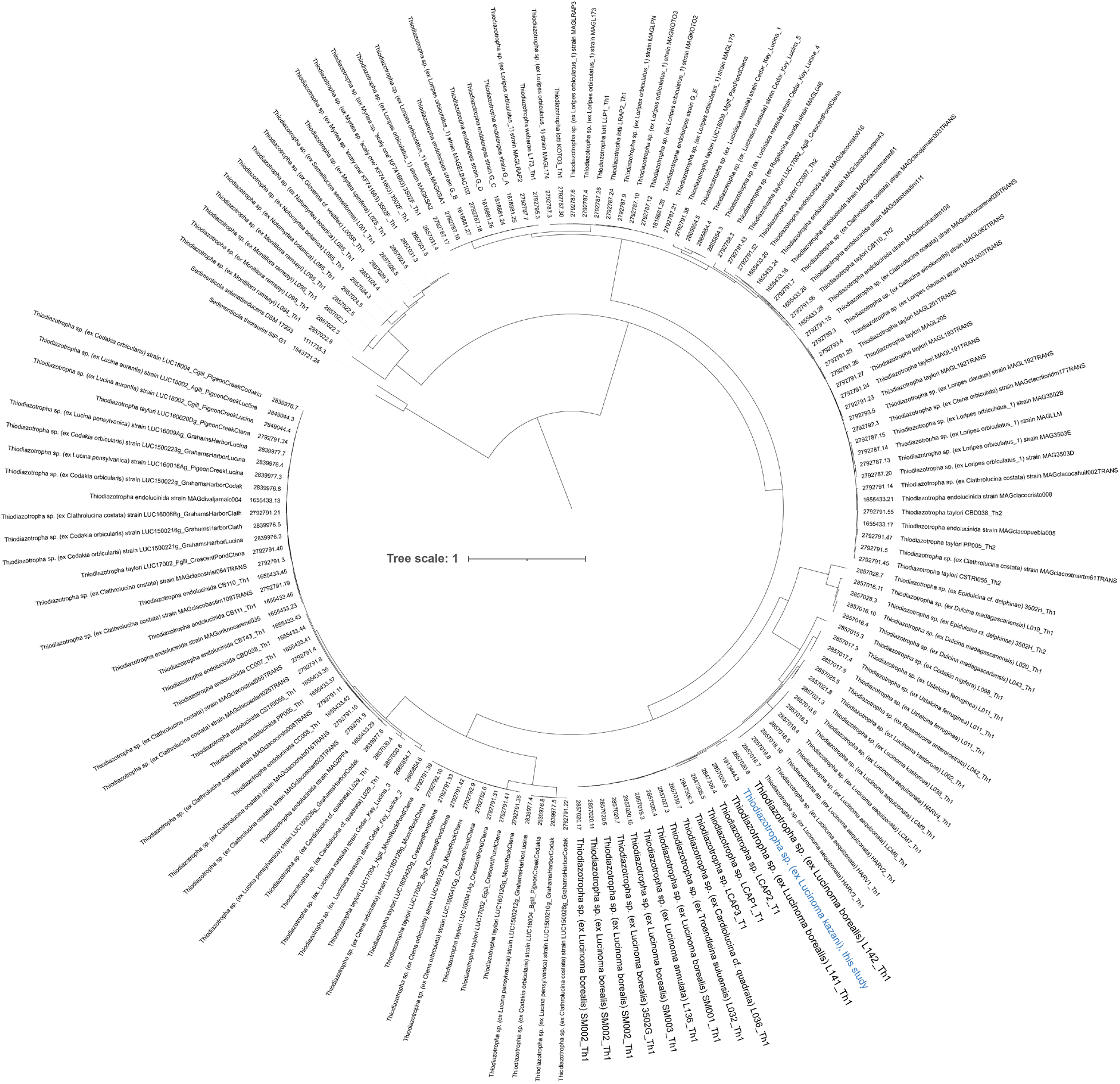
Phylohgenomic tree of 150 genomes in the Thiodiazotropha clade (2 *Sedimenticola* genomes were added for an outgroup). The tree is based on 411 single-copy genes and uses the JTT model in RAxML. The tree is drawn to scale and rooted at the midpoint. Numbers represent BV-BRC accessions.

**Supplementary Figure S2:**
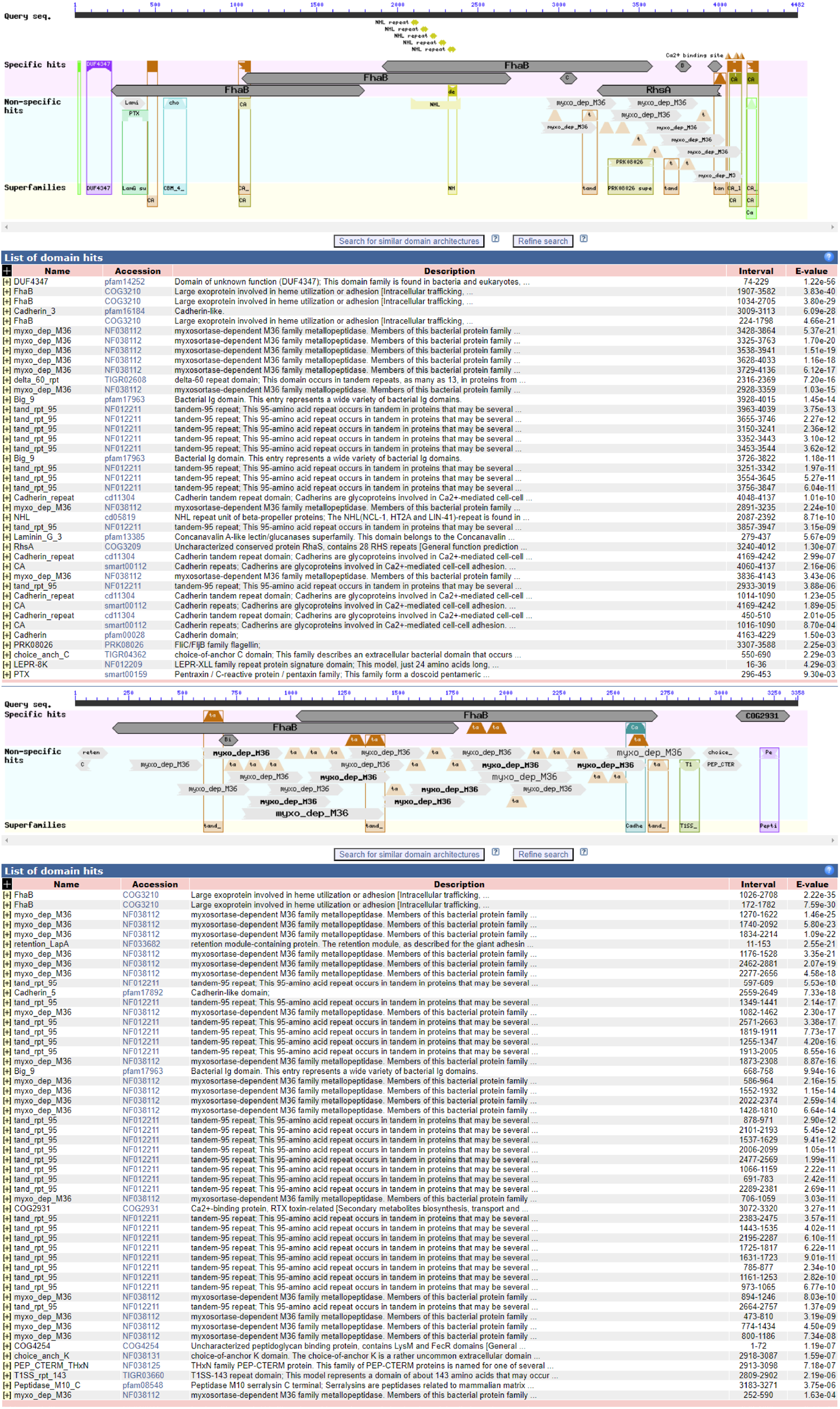
Domains in the trwo large toxin-like proteins in *Lucinoma kazani* genomes (CD-search, NCBI).

**Supplementary Figure S3:**
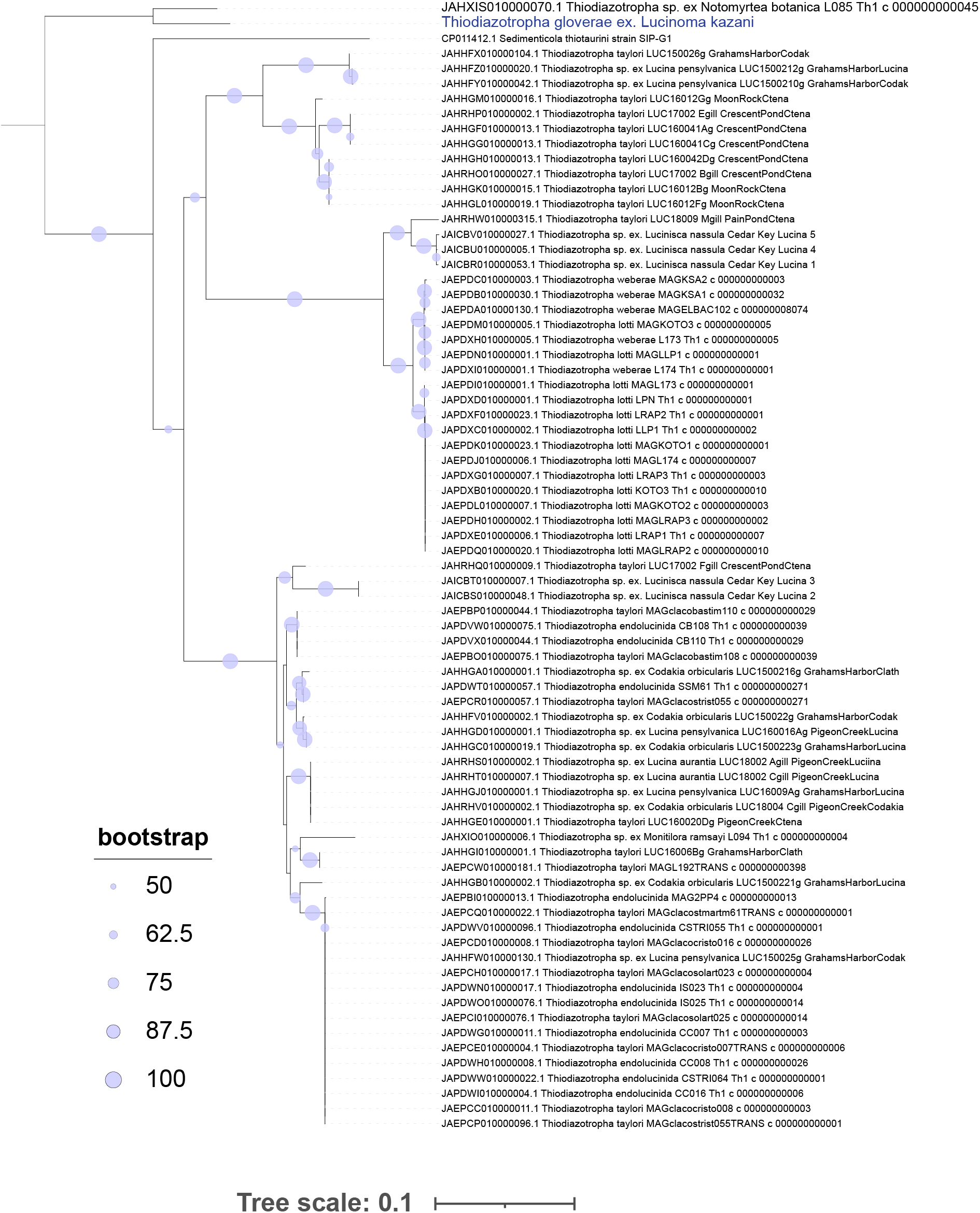
Phylogeny of 73 nucleotide sequences encoding NifD in the Thiodiazotropha clade. The maximum likelihood tree is constructed in IQtree using the TN+F+I+G4 model. The tree is rooted at the midpoint. The tree scale represents the number of substitutions per site.

### Supplementary Table Legends

**Supplementary Table ST1**: Stable isotope analyses of carbon, nitrogen and sulfur in *Lucinoma kazani* tissues.

**Supplementary Table ST2**: Genomic features of *Ca.* Thiodoazotropha ex. *Lucinoma kazani*. Expression values as RNA read counts and protein abundance as % normalized spectral abundance factor (NSAF) in gills samples of four *L. kazani* specimens. Red fonts represent annotations based on homology with NCBI/UniProt databases (defined as hypotheticals by RAST-tk annotation). Green cells represent putative prophage regions.

**Supplementary Table ST3**: Protein annotations, Figure 2.

